# Characterization of the *Pristionchus pacificus* “epigenetic toolkit” reveals the evolutionary loss of the histone methyltransferase complex PRC2

**DOI:** 10.1101/2023.12.05.570140

**Authors:** Audrey Brown, Adriaan B. Meiborg, Mirita Franz-Wachtel, Boris Macek, Spencer Gordon, Ofer Rog, Cameron J Weadick, Michael S. Werner

## Abstract

Comparative approaches have revealed both divergent and convergent paths to achieving shared developmental outcomes. Thus, only through assembling multiple case studies can we understand biological principles. Yet, despite appreciating the conservation – or lack thereof – of developmental networks, the conservation of epigenetic mechanisms regulating these networks is poorly understood. The nematode *Pristionchus pacificus* has emerged as a model system of plasticity and epigenetic regulation as it exhibits a bacterivorous or omnivorous morph depending on its environment. Here, we determined the “epigenetic toolkit” available to *P. pacificus* as a resource for future functional work on plasticity, and as a comparison with *C. elegans* to investigate the conservation of epigenetic mechanisms. Broadly, we observed a similar cast of genes with putative epigenetic function between *C. elegans* and *P. pacificus*. However, we also found striking differences. Most notably, the histone methyltransferase complex PRC2 appears to be missing in *P. pacificus.* We described the deletion/pseudogenization of the PRC2 genes *mes-2* and *mes-6* and concluded that both were lost in the last common ancestor of *P. pacificus* and a related species *P. arcanus.* Interestingly, we observed the enzymatic product of PRC2 (H3K27me3) by mass spectrometry and immunofluorescence, suggesting that a currently unknown methyltransferase has been co-opted for heterochromatin silencing. Altogether, we have provided an inventory of epigenetic genes in *P. pacificus* to enable reverse-genetic experiments related to plasticity, and in doing so have described the first loss of PRC2 in a multicellular organism.

## Introduction

Comparative studies have revealed how developmental networks can evolve over time; sometimes generating the same phenotype, and sometimes leading to novel traits (Peel 2008; Peter and Davidson 2011). These studies have primarily focused on the conservation of genes underlying trait development. Meanwhile, comparatively little attention has been paid to the evolution of factors that regulate development. Though the term “epigenetic” has taken on various meanings over time, the current definition refers to mitotically and/or meiotically stable, non-DNA sequence based mechanisms that regulate gene expression (Jablonka and Lamb 2002). Epigenetic mechanisms can be divided into three categories: DNA modification (e.g., 5mC), histone post-translational modification (e.g., H3K27me3), and non-coding RNA pathways (e.g., RNAi) (Liebers et al. 2014; Allis and Jenuwein 2016). Studying the evolution of epigenetic pathways may offer insights into developmental robustness vs. plasticity and the evolution of novel traits (Levis and Pfennig 2019).

The nematode *Pristionchus pacificus* was established in the 1990’s as a comparative system to the model nematode *Caenorhabditis elegans* (last sharing a common ancestor 80-200 mya) (Howard et al. 2022). Having two divergent, experimentally tractable model organisms in the same Order has facilitated functional evolutionary and developmental (evo-devo) studies leading to several unexpected findings (Markov et al. 2016; Rošić et al. 2018; Beltran et al. 2019; Bui and Ragsdale 2019; Hong et al. 2019; Weadick 2020; Sun et al. 2021; Lo et al. 2022). For example, early studies on vulva development revealed the extent of developmental systems drift, whereby the genetic underpinnings of a trait diverge even as the trait itself is conserved (Wang and Sommer 2011; Sommer 2012). However, a thorough exploration of the conservation of epigenetic pathways between the two species has not yet been done.

There is also interest in characterizing the epigenetic pathways in *P. pacificus* through the lens of developmental plasticity, a phenomenon where environmental conditions experienced during development influence adult phenotypes. *P. pacificus* displays morphological plasticity of its feeding structures: adult worms exhibit either an omnivorous or bacterivorous mouth-form depending on signals experienced as juveniles (Bento et al. 2010). The difference between these forms is multifaceted. The microbivorous morph is narrow, deep, and contains a single dorsal “tooth” while the omnivorous morph is wide, shallow, and contains two teeth. Several studies have now pointed to epigenetic mechanisms as a way to translate environmental signals into gene expression changes that underly the development of different traits (Valena and Moczek 2012; Duncan et al. 2014; Yan et al. 2014; Kilvitis et al. 2017; Thorson et al. 2017; Budd et al. 2022; Toker et al. 2022; Jordan et al. 2023; Werner et al. 2023). We recently showed that perturbing histone acetylation/deacetylation with chemical inhibitors alters both mouth-form phenotype and the expression of key “switch genes,” indicating that epigenetic pathways are, at least in part, responsible for the regulation of mouth-form development (Werner et al. 2023). However, further genetic and biochemical experiments are needed to reveal how histone modifications regulate plasticity in this system, which requires prior knowledge of the of the “writers” and “erasers” of modifications present in *P. pacificus*. In essence, in order to manipulate proteins that effect epigenetic regulation, we first need to know which those proteins are.

As a relatively new model system, *P. pacificus* lacks a well characterized “toolkit” of genes and modifications with epigenetic function. A high-quality chromosome-scale genome assembly with manually curated gene annotations has been created for *P. pacificus* (Rödelsperger et al. 2017; Athanasouli et al. 2020), yet the annotations lack thorough functional categorization compared to other model systems backed by decades of experimental research. To address this gap, we identified putative *P. pacificus* epigenetic genes using an orthology and domain-informed annotation pipeline. We then used LC-MS/MS to identify the histone post-translational modifications (PTMs) present in *P. pacificus* and, by orthology, predict which proteins may have added or removed these observed marks. Surprisingly, after curating this inventory we found that the highly conserved methyltransferase PRC2 (Polycomb Repressive Complex 2) was missing in *P. pacificus* despite the presence of its associated mark, H3K27me3. We investigated the presence/absence of the PRC2 complex in the *Pristionchus* phylogeny and traced the loss of two of its components (MES-2 and MES-6) to the last common ancestor of *P. pacificus* and its relative *P. arcanus.* To our knowledge, its absence in *P. pacificus* represents the first identified loss in a multicellular organism. The presence of H3K27me3 in the absence of PRC2 indicates that a currently unknown methyltransferase “writes” H3K27me3 and maintains its role in gene silencing. In summary, we produced a dataset of the “epigenetic toolkit” available to *P. pacificus* as a resource for future comparative studies with *C. elegans* and experimental work in developmental plasticity; and revealed the evolutionary loss of a highly conserved methyltransferase complex.

## Methods

### Identification of putative epigenetic genes

We used a domain and orthology-informed pipeline adapted from Pratx et al. 2018 to predict genes in *P. pacificus* and *C. elegans* with putative epigenetic function. As input, we used the proteomes of 17 species, including six well-studied model organisms (*Caenorhabditis elegans*, *Drosophila melanogaster*, *Homo sapiens, Arabidopsis thaliana, Saccharomyces cerevisiae*, *Schyzosaccharomyces pombe)*, two emerging model systems for developmental plasticity (*Acyrthosiphon pisum, Apis mellifera*), six additional nematodes (*Pristionchus pacificus*, *Pristionchus exspectatus*, *Pristionchus mayeri*, *Brugia malayi*, *Strongyloides ratti*, *Trichinella spiralis*), plus three additional species — an animal (*Danio rerio*), fungus (*Leptosphaeria maculans)* and plant (*Selaginella moellendorffii)* — included to help improve phylogenetic resolution. Proteome sources and accession numbers are indicated in Supplementary Table 1 (Athanasouli et al. 2020; Rödelsperger 2021; UniProt Consortium 2021).

To identify putative epigenetic genes in our target species (*P. pacificus*), we relied on reference datasets of known epigenetic proteins and epigenetic-associated protein domains (i.e., a protein or domain previously shown to be exclusively linked to epigenetic processes) in the six model species included in our above input (*C. elegans, H. sapiens, A. thaliana, S. cerevisiae, S. pombe,* and *D. melanogaster*). We initially obtained these reference datasets from Pratx *et al*. (2018) However, we adjusted them to reflect updated UniProt accession numbers and to include an additional 54 epigenetic proteins that we identified from UniProt and additional literature and database searches on top of the original 691 from Pratx et al. 2018 (Additional Files 1-2).

The pipeline consists of six steps (Fig. 1):

1. A Pfam domain search on all *P. pacificus* proteins using InterProScan software, version 5.58-91.0, with default search settings (Additional File 3; Jones et al. 2014).
2. Isolation of *P. pacificus* proteins containing epigenetic-associated Pfam domains (based on the reference epigenetic-associated protein domain dataset).
3. Orthology clustering of the 17 proteomes using OrthoFinder version 2.5.4 software, default settings (Emms and Kelly 2019). From OrthoFinder, we obtained “orthogroups” (i.e., groups of orthologous proteins) and orthogroup gene phylogenies (Additional File 4).
4. Isolation of *P. pacificus* proteins from orthogroups containing one or more known epigenetic proteins (based on the reference model organism epigenetic protein dataset).
5. Quality control to remove false positives, including validating orthogroup phylogenies and orthogroup domain composition, and reannotation of gene annotation errors (Additional File 5). Additional details on quality control steps are provided below.
6. Merging the orthology-identified and domain-identified *P. pacificus* protein datasets and assembly of a final epigenetic gene dataset from the genes encoding each protein. This was done to account for cases where multiple genes encode the same protein sequence (i.e., duplicate genes), or where a single gene produces multiple isoforms (often assigned distinct UniProt entries). Epigenetic genes were manually sorted into functionally defined families (i.e., histone acetyltransferases, histone deacetylases, etc.) based on previously characterized orthology and domain predictions (Additional File 6).

**Figure 1:**
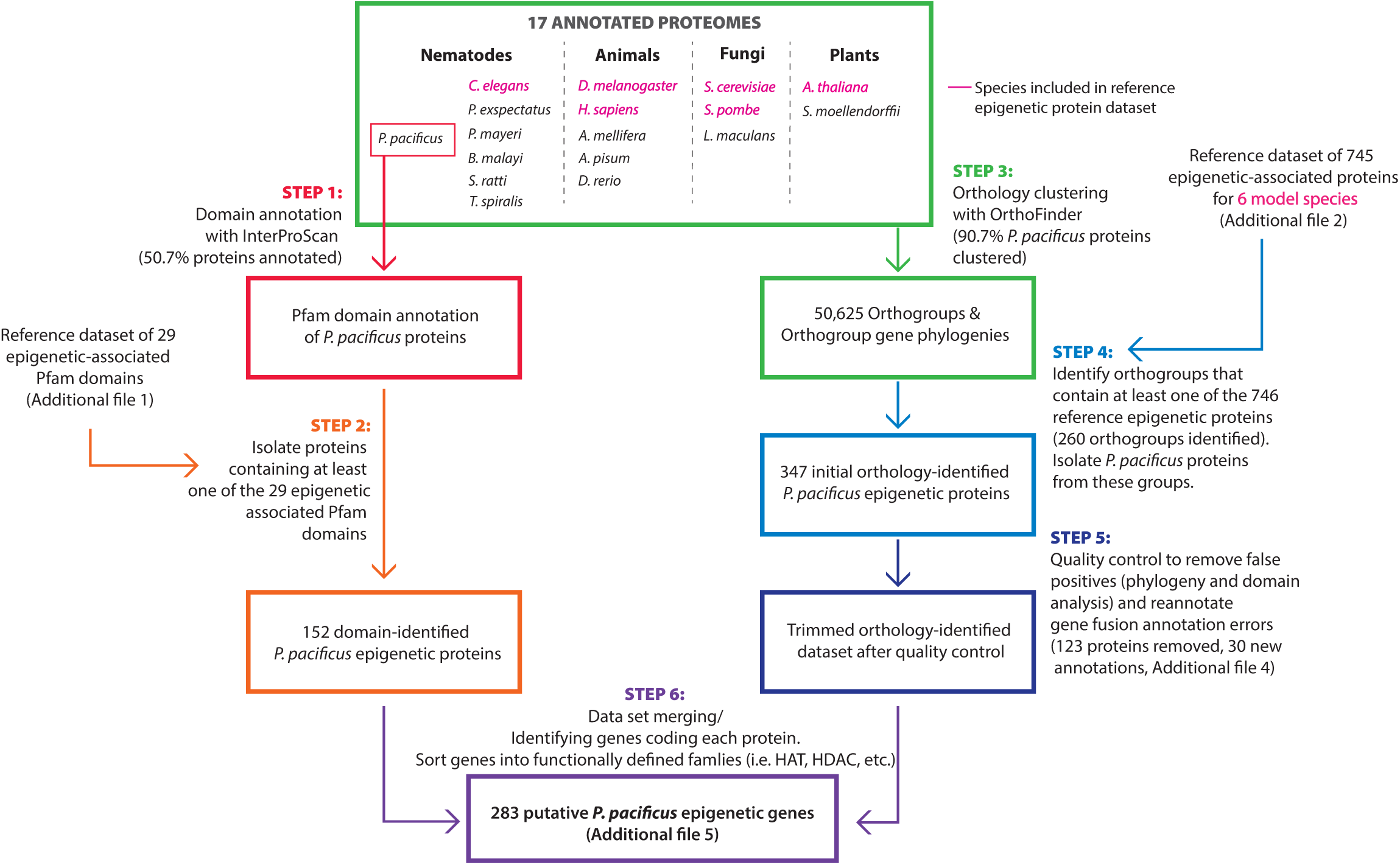
Pipeline for identifying putative epigenetic genes in *P. pacificus*. (1) *P. pacificus* Pfam protein domains were annotated using lnterProScan. (2) *P. pacificus* proteins containing at least one epigenetic-associated Pfam domain were identified. (3) All proteins from *P. pacificus* and 16 other species were clustered into orthogroups using OrthoFinder. (4) Orthogroups containing known epigenetic proteins from at least one of 6 reference model species were identified, and *P. pacificus* proteins from these were isolated. (5) This orthology-identified dataset was subjected to a series of quality control measures: analyzing orthogroups for phylogenetic correctness and consistent domain composition, and manual re-annotation of gene fusion annotation errors. (6) The domain-identified and orthology-identified datasets were merged, and protein-coding genes were compiled into a final epigenetic gene dataset.

Prior to any filtering, 91.8% of the 28,896 proteins included in the *P. pacificus* annotation were either sorted into an OrthoFinder orthogroup or annotated with a Pfam domain by InterProScan (irrespective of annotated function); the remaining 8.2% presumably reflect lineage-specific genes and/or poor annotations. To validate our pipeline and to produce a comparative dataset to *P. pacificus,* the same pipeline was used to identify *C. elegans* epigenetic genes, with the modification of first removing *C. elegans* proteins from the reference model organism epigenetic protein dataset (Additional Files 7-9).

### Orthology quality control

In step #5 of our pipeline, we performed three quality control measures:

1. We manually checked the placement of *P. pacificus* genes for phylogenetic correctness within each orthogroup by viewing the resolved gene tree output from OrthoFinder with Dendroscope version 3.8.3 (Huson and Scornavacca 2012). Unless they contained an epigenetic-specific protein domain, *P. pacificus* genes were removed from the dataset if they 1) were a member of a tree or subtree that did not broadly recapitulate known phylogenetic relationships, or 2) were phylogenetically misplaced in an otherwise acceptable tree/subtree. Given that the gene trees were based on single gene alignments, and considering the broad phylogenetic scale involved, we did not require perfect recapitulation of established relationships; rather, we simply checked for clear and obvious departures suggestive of orthology group errors.
2. We performed a domain search on each protein within the candidate orthogroups using InterProScan and manually validated these with NCBI’s CD-Search (Lu et al. 2020). For each orthogroup, we identified *P. pacificus* proteins with “outlier” domains that were not found in any other protein within the orthogroup. Such proteins were discarded if they also 1) did not contain any epigenetic-specific domains, 2) had no predicted epigenetic functions (from NCBI’s CD-Search), or 3) were not closely related to other *P. pacificus* or model organism epigenetic proteins (using the phylogenies from #1). In addition, we also excluded proteins with no predicted domains unless they were close orthologs of another *P. pacificus* or model organism epigenetic protein.
3. We manually reannotated several gene fusion annotation errors in *P. pacificus* histone genes. These were cases where we detected the presence of multiple core histone domains within single gene annotation entries. Manual inspection of start/stop codon positions and existing Iso-seq reads (Werner et al. 2018) revealed these were in fact distinct histone genes, closely linked in tandem (Supplementary Fig. 6, Additional File 5).

### Statistics on epigenetic gene counts

To compare the gene counts per epigenetic category between *P. pacificus* and *C. elegans,* we performed Fisher’s exact tests using Monte Carlo simulation on the *C. elegans* and *P. pacificus* putative epigenetic gene count datasets (Table 1). This was done first on the protein family-level sums (8×2) and next on the sub-family-level sums (33×2). We also performed 2×2 Fisher’s exact tests on contingency tables of each protein family-level sum vs. the sum of all others, with the p-value significance threshold adjusted via Bonferroni correction to account for multiple testing.

**Table 1:** Protein coding epigenetic genes in *P. pacificus* and *C. elegans.* Table summary of epigenetic genes identified in *P. pacificus* and *C. elegans*. Proteins were classified into families and subfamilies based on domain composition. P-values were calculated using Fisher’s exact tests on 2×2 contingency tables of *P. pacificus* and *C. elegans* protein family sums vs. the sum of the remaining epigenetic genes. *p<0.05 (remains significant after Bonferroni correction).

| Family | Subfamily | *P. pacificus* | *C. elegans* | P |
| --- | --- | --- | --- | --- |
| HAT | CBP/P300 | 12 | 7 | 0.170 |
|  | GNAT | 15 | 11 |  |
|  | MYST | 7 | 4 |  |
|  | Other | 2 | 1 |  |
| HDAC | Class I | 4 | 3 | 1 |
|  | Class II | 4 | 5 |  |
|  | Class III | 5 | 4 |  |
|  | Class IV | 1 | 1 |  |
| HMT | PRMT | 12 | 3 | 0.064 |
|  | SET | 45 | 31 |  |
|  | DOT1 | 6 | 6 |  |
|  | Other | 1 | 1 |  |
| HDMT | JMJ | 20 | 12 | 0.399 |
|  | LSD | 2 | 3 |  |
| Histones | H1 | 5 | 9 | 0.004* |
|  | H2A | 13 | 19 |  |
|  | H2B | 12 | 17 |  |
|  | H3 | 19 | 24 |  |
|  | H4 | 12 | 16 |  |
| DNA methylation | DNMT | 1 | 0 | 1 |
|  | ALKB | 5 | 5 |  |
|  | DAMT-1 | 1 | 1 |  |
| ncRNA associated | Dicer | 1 | 1 | 0.609 |
|  | RdRP | 5 | 4 |  |
|  | Helicases | 7 | 4 |  |
|  | Argonautes | 23 | 20 |  |
|  | Drosha | 1 | 1 |  |
|  | Microprocessor complex | 2 | 1 |  |
| Others | HK | 17 | 22 | 0.406 |
|  | HP | 5 | 6 |  |
|  | HUT | 9 | 7 |  |
|  | H DU | 6 | 6 |  |
|  | Misc. | 3 | 2 |  |
| Total |  | 283 | 257 |  |

### Sequence alignments and phylogenetic construction

All protein and nucleotide sequence alignments were created using the online version of MAFFT version 7 (Katoh et al. 2019). Orthogroup phylogenies were retrieved from OrthoFinder as described above. To construct all other phylogenies, protein sequence alignments were processed with ClipKIT (Steenwyk et al. 2020), and phylogenies were constructed using IQ-TREE 2 with the automated model testing setting and 1000 bootstrap replicates (Minh et al. 2020). Phylogenies were graphed using either the R package “ggtree” (unrooted trees) or “phangorn” (midpoint rooted trees) (Schliep 2011; Yu et al. 2017). All tools were run using default settings unless otherwise indicated.

### Histone categorization and mapping

1. *C. elegans* canonical and variant H2A, H2B, H3, and H4 histone sequences have been previously characterized (Pettitt et al. 2002; Keall et al. 2007). We identified the *P. pacificus* canonical histone sequences as those genes with the best TBLASTN hit using the *C. elegans* canonical sequences as query-all other sequences were designated as variants. 95% of *P. pacificus* and 98% of *C. elegans* canonical genes also contained a conserved hairpin sequence in the 3’ UTR, indicating they are replication-dependent (RD) (Supplementary Fig. 2B; Pettitt et al. 2002). Nucleotide sequence logos of the RD 3’ UTR hairpin sequence were plotted from nucleotide alignments of 100 bases immediately 3’ of all the stop codons of canonical histones using the “ggseqlogo” R package (Wagih 2017). Histone chromosomal positions were mapped using custom code written in R version 4.4.2 (https://github.com/audreybrown1/Brown-et-al.-2023-library, Additional File 13-14). Histone clusters in *C. bovis, P. mayeri,* and *A. sudhausi* were identified by BLASTP of H4 as it is the most conserved histone with the fewest variants, and then manually searching for other histone genes in the vicinity. Note, the genomes of *P. mayeri* and *A. sudhausi* are assembled from short-read Illumina sequences and are of poorer overall quality; thus we are likely missing several histone gene clusters in these species. All genetic resources used are indicated in Supplementary Table 1.

### Histone Liquid Chromatography Tandem-Mass Spectrometry (LC-MS/MS)

Histones from *P. pacificus* were acid-extracted following Werner et al. 2023. Briefly, cultures of *P. pacificus* were synchronized by bleach-NaOH, and eggs were aliquoted on to standard NGM-agar plates. Worm pellets (∼200-500 µl size) were collected and flash-frozen in liquid N2 after 72 hours, representing primarily adults. Crude nuclei preparations were made from worm pellets and then histones were extracted using sulfuric acid following Schecter et al. 2007 (Shechter et al. 2007). After extraction, 50-100 µg of histones were digested with Arg-C following reduction and alkylation. To stop the digestion reaction, 10% trifluoroacetic acid was added to a final concentration of 0.5%.

Analysis of digested histones was done on an Easy-nLC 1200 system coupled to a QExactive HF-X mass spectrometer (both Thermo Fisher Scientific) as described in Werner et al. 2023. MS data were analyzed by MaxQuant software version 1.5.2.8 (Cox and Mann 2008) with integrated Andromeda search engine (Cox et al. 2011). Endoprotease ArgC was defined as protease with a maximum of two and three missed cleavages, respectively. The minimum peptide length was set to five. Carbamidomethylation on cysteine was set as fixed modification in all processings of different combinations of variable modifications. These were mono-, di- and trimethylation on lysine and arginine, phosphorylation on serine, threonine and tyrosine, acetylation, crotonylation, butyrylation, hydroxybutyrylation, propionylation, and di-glycine on lysine, and O-GlcNacylation on serine and threonine. No more than five modifications were searched at the same time.

Data was mapped to the *P. pacifius* ‘El Paco’ protein annotation version 1 (Rödelsperger et al. 2017), with a quality control threshold score >100 and posterior error probability (PEP) <0.01. Initial maximum allowed mass tolerance was set to 4.5 parts per million (ppm) for precursor ions and 20 ppm for fragment ions. Peptide, protein and modification site identifications were reported at a false discovery rate (FDR) of 0.01, estimated by the target/decoy approach (Elias and Gygi 2007).

### Prediction of *P. pacificus* histone PTM writers

*P. pacificus* orthologs of previously characterized *C. elegans* and human histone-modifying proteins were identified based on manual inspection of their previously generated orthogroup assignments and the associated orthogroup phylogenies. Human and *C. elegans* proteins known to “write” or “erase” specific acetyl and methyl modifications were identified from literature and database searches. For brevity, we focused on acetyltransferases, deacetylases, methyltransferases and demethylases (Additional File 9).

### PRC2 ortholog identification in *Pristionchus* nematodes

We used the genomes, protein sequences, and gene annotations of ten diplogastrid nematode species to identify and analyze the conservation of the PRC2 complex within the *Pristionchus* lineage: *Pristionchus pacificus, Pristionchus exspectatus, Pristionchus arcanus, Pristionchus maxplancki, Pristionchus japonicus, Pristionchus mayeri, Pristionchus entomophagus, Pristionchus fissidentatus, Micoletzkya japonica,* and *Parapristionchus giblindavisi.* All genetic resources used for these species, including genome assemblies, transcriptomes, and RNA sequencing reads, are indicated in Supplementary Table 1. Genomes were viewed using IGV version 2.12.2 (Robinson et al. 2011).

We used BLAST and OrthoFinder to search for orthologs of the PRC2 proteins MES-2/EZH2 (*C. elegans* ID/Human ID), MES-3/SUZ12, and MES-6/EED in these ten species. For BLAST, we used BLASTP to search predicted proteomes and TBLASTN to search the assembled transcriptomes, and genomes (and also in *P. pacificus,* the raw PacBio reads) of each nematode species, using both the human and *C. elegans* PRC2 sequences as query. A significance threshold for BLASTP and TBLASTN of the transcriptome was set by plotting the e-values of the top hits (pooled across all ten species) and then identifying where the hit e-values plateaued: based on this approach, we chose a threshold of 10e-29 (Supplementary Fig. 5A-B). As expected, TBLASTN to the whole genome/raw PacBio reads produced fewer and lower significance hits (results not shown), and therefore, a threshold of 10e-5 was used. For OrthoFinder, we clustered the above-listed ten species plus *C. elegans* and humans to search for PRC2 orthologs of human and *C. elegans* PRC2 components in each of the ten diplogastrid nematodes.

We also manually searched for *mes-2* and *mes-6* orthologs, and confirmed the annotations of previously identified orthologs, using formerly generated RNA-seq reads and positional information. For species in which we found an ortholog using BLAST or OrthoFinder, we isolated the genomic region spanning approximately 50,000-70,000 bases upstream and downstream and identified neighboring genes. Then, for species in which a *mes-2/mes-6* ortholog could not be found using BLAST or OrthoFinder, we searched for orthologs of neighboring genes to isolate the same genomic region. Each region was visually inspected in IGV alongside mixed-stage RNA-seq data (Rödelsperger et al. 2018) that was mapped to the genomes using the STAR sequence alignment software, version 2.7.10 (Dobin et al. 2013). Newly identified orthologs and gene annotations that did not match the RNA transcript data were manually re-annotated (Additional File 5). Additionally, we used previously annotated repetitive sequences for *P. pacificus* to search the focal genomic regions for the presence of repetitive sequences indicative of transposons (Athanasouli and Rödelsperger 2022).

### Synteny analysis

We arranged each genomic region from the previous analysis based on the *Pristionchus* phylogeny (we excluded *M. japonica* and *P. giblindavisi* from this analysis due to short scaffold lengths). For each gene within this region, start and stop nucleotide positions were plotted using custom R code (https://github.com/audreybrown1/Brown-et-al.-2023-library). Genes were grouped according to their previously calculated orthogroups (Additional File 11-12).

### RNA-seq analysis

To analyze the expression of the PRC2 genes *mes-2* and *mes-6* across species, we obtained mixed stage, paired-end RNA-seq data for *P. fissidentatus* (1 replicate)*, P. entomophagus* (1 replicate)*, P. mayeri* (1 replicate)*, P. japonicus* (1 replicate)*, P. maxplancki* (1 replicate)*, P. arcanus* (5 replicates)*, P. exspectatus* (5 replicates), and *P. pacificus* (5 replicates) from the European Nucleotide Archive (accession number PRJEB20959; Rödelsperger et al. 2018). Reads were aligned to the reference genomes using the STAR version 2.7.10 sequence alignment software, SAM output files were processed using Samtools version 1.16, and reads were quantified using htSeq-count version 2.0.2 with the nonunique none setting (Dobin et al. 2013; Anders et al. 2015; Danecek et al. 2021). Expression for each gene was quantified as FPKM:

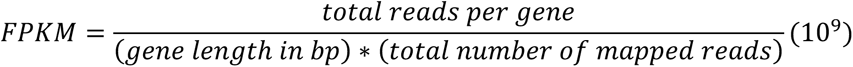

### Immunofluorescence

Immunofluorescence was performed as previously described (Phillips et al. 2009). In brief, adult *P. pacificus* or *C. elegans* were picked into Egg buffer with 0.1% Tween and 0.06% Sodium Azide, dissected, and fixed with Egg buffer and 1% formaldehyde. Dissected worms were transferred onto HistoBond slides and frozen on dry ice for >1 minute. The coverslip of frozen samples was quickly removed, and worms were washed in -20°C methanol for 1 minute, then washed for 5 minutes 3x in room temperature PBS-Tween. Worms were blocked in Roche Block solution for 30 minutes and then incubated overnight in a 1:500 or 1:1000 dilution of primary antibodies. After incubation, worms were washed for 10 minutes 3x with PBS-Tween and then incubated in a 1:500 secondary antibody dilution for 1 ½ hours. Following secondary antibody incubation, worms were washed for 10 minutes each in PBS-Tween, PBS-Tween with 0.5 mg/ml DAPI, and again with PBS-Tween. Slides were mounted with N-propyl gallate-glycerol. Confocal images were taken using a Zeiss LSM 880 confocal microscope equipped with an AiryScan and using a 63x 1.4 NA oil immersion objective. Maximum intensity projection confocal images are shown throughout, and were processed using Zen Blue 3.0 and ImageJ. Primary antibodies used were Rabbit Tri-Methyl-Histone H3 (C36B11) (Cell Signaling, #9733) and Histone H3 (1B1B2) Mouse mAb (Cell Signaling, #14269). Secondary antibodies used were Goat bAb to Rb IgG Alexa Fluor 647 (Abcam, #150079) and Goat anti-Mouse IgG3 Cross-Adsorbed Secondary Antibody Alexa Fluor 594 (ThermoFisher, # A21155).

## Results

### An inventory of *P. pacificus* and *C. elegans* epigenetic genes

We identified the “epigenetic toolkit” available to *Pristionchus pacificus* through a combination of orthology and domain predictions similar to an approach by Pratx et al. 2018 (Fig. 1). Broadly, proteins and protein domains that are linked to epigenetic processes in a diverse set of model systems were used to identify orthologs in our focal study system. Specifically, we assessed whether each *P. pacificus* protein (1) has domains that are found exclusively in epigenetic-function associated proteins and/or (2) is orthologous to proteins in model species with known epigenetic function. To make these assessments, we relied on reference datasets of proteins connected to DNA modification, histone modification, or non-coding RNA pathways in six model species (Additional Files 1-2). These were initially curated by Pratx et al., 2018, but were updated with an additional 54 proteins identified from further literature and database searches, for a total of 745 reference epigenetically-associated proteins and 29 reference epigenetically-associated domains. This approach allowed us to capture orthologs in a high-throughput manner and identify genes that may be otherwise missed in 1:1 BLAST comparisons due to gene loss, gain or divergence.

A total of 260 orthogroups contained reference model species epigenetic proteins, and 126 of these contained one or more *P. pacificus* proteins. From these groups, an initial 347 putative *P. pacificus* epigenetic proteins were identified. Manual curation of this list resulted in 123 proteins being filtered out from our dataset (see methods). For example, orthogroups containing epigenetically associated kinases tended to be extremely large and often contained many other proteins with non-epigenetic functions. As a result, we only included kinases where a close phylogenetic relationship with a known model-organism histone kinase could be observed. After quality control, we retained 96 orthogroups with epigenetic-function that contained a *P. pacificus* gene, 40 of which contained more than one *P. pacificus* gene: 5 orthogroups contained *P. pacificus-*specific expansions, 28 contained deeper, pre-speciation expansions, and 7 orthogroups contained both. Ultimately, we produced a final dataset of 283 *P. pacificus* epigenetic-function genes (Table 1, Additional File 6).

We also annotated putative *C. elegans* epigenetic genes using our pipeline to produce a comparative dataset (but with the prior removal of all *C. elegans* genes from our reference datasets; 156 total genes, 101 unique protein sequences). Here, our pipeline and quality control yielded a final dataset of 257 *C. elegans* epigenetic-associated genes (Table 1). We recovered 152 of the 156 (97%) reference genes we had removed before applying the pipeline, validating it’s efficiency. The four unrecovered reference genes each belong to a *C. elegans-*specific orthogroup and lack distinct epigenetic domains, which is presumably why we could not identify them from our approach. We also identified 105 additional putative epigenetic genes not previously included in the *C. elegans* reference set. Upon closer inspection, 47 were previously annotated in UniProt as having epigenetic functions (relating to DNA methylation, histone modification, or RNAi). Therefore, these likely represent bona fide epigenetic genes that were absent in our manually updated reference dataset. The remaining 58 had no explicit epigenetic annotations and are likely the result of duplication and divergence of other epigenetic-associated genes. Taken together, this analysis suggests that 1) our pipeline is more effective at identifying epigenetic proteins in well-annotated organisms than literature or database searches and 2) our pipeline can identify “new” putative epigenetic proteins, even in organisms with decades of functional characterization.

Next, we compared the two focal species nematode data sets. We manually sorted genes from the *P. pacificus* and *C elegans* datasets into protein families based on their domain or subfamily composition (Table 1). Phylogenetic analysis of these families shows that within each, there are several instances of one-to-one orthology between *P. pacificus* and *C. elegans* (Supplementary Fig. 1). However, we also observed multiple instances where proteins have no ortholog within the other species, confirming that the inclusion of species beyond *C. elegans* in our bioinformatic pipeline was necessary to identify such proteins (Supplementary Fig. 1). Comparing the size of the gene families (i.e., Fisher’s exact test on the 8×2 contingency table of family-by-species sums) suggested there might be a difference in specific epigenetic pathways (p= 0.065). However, we found no evidence for a difference between the two species at the subfamily level (p=0.941, Fisher’s exact test on the 33×2 contingency table of subfamily-by-species sums). To further explore differences at the gene family level, we compared the size of each individual family against the sum of all other genes. This analysis indicated a significant difference in the number of histone genes between species: *C. elegans* has a disproportionately large histone complement compared to *P. pacificus* (p=0.004; Table 1). Indeed, this pattern appears to be consistent across each histone subfamily — *C. elegans* contains more copies of H1, H2A, H2B, H3 and H4 than *P. pacificus*.

### *P. pacificus* and *C. elegans* histone gene clusters are deeply diverged

In metazoans, histone genes have repeatedly undergone serial duplication events, and duplicate copies of H2A, H2B, H3, and H4 histones can be found clustered in groups, and distributed throughout the genome many times over (Amatori et al. 2021). These clusters tend to encode “canonical” histones: canonical histones are nearly identical at the amino acid level and are replication-dependent (RD), with expression restricted to S-phase when the nucleosomes are doubled after DNA replication (Amatori et al. 2021). By contrast, “variant” histones often reside ouside of these repeated arrays, have slightly modified amino acid sequences, and are expressed throughout the cell cycle. After finding that *C. elegans* contains a larger histone gene compliment than *P. pacificus,* we asked whether differences in the number of canonical or variant histones drove this variation. To answer this question, we examined the differences in canonical and variant histones between the two species using unrooted phylogenies of all *P. pacificus* and *C. elegans* histones generated from amino acid sequence alignments (Fig. 2A-D, Supplementary Fig. 2C). Within the core nucleosome histones (H2A, H2B, H3, and H4), *C. elegans* has 61 canonical genes and 15 variants, while *P. pacificus* has 41 canonical and 15 variants. Therefore, canonical histones appear to be driving the difference in histone gene number between the two species rather than histone variants. Answering this question also allowed us to observe additional differences between *C. elegans* and *P. pacificus* canonical histone sequences. First, we observed that *P. pacificus’* and *C. elegans’* canonical histones (except H4) differ slightly in amino acid sequence. H2A differs by seven amino acids, H3 by three amino acids, and H2B by 12-30 amino acids (*C. elegans* contains multiple canonical H2B sequences; Supplementary Fig. 2A). This follows a known pattern that H2A and H2B are more labile at the amino acid sequence level than H3 and H4 (Thatcher and Gorovsky 1994; Raman et al. 2022).

**Figure 2:**
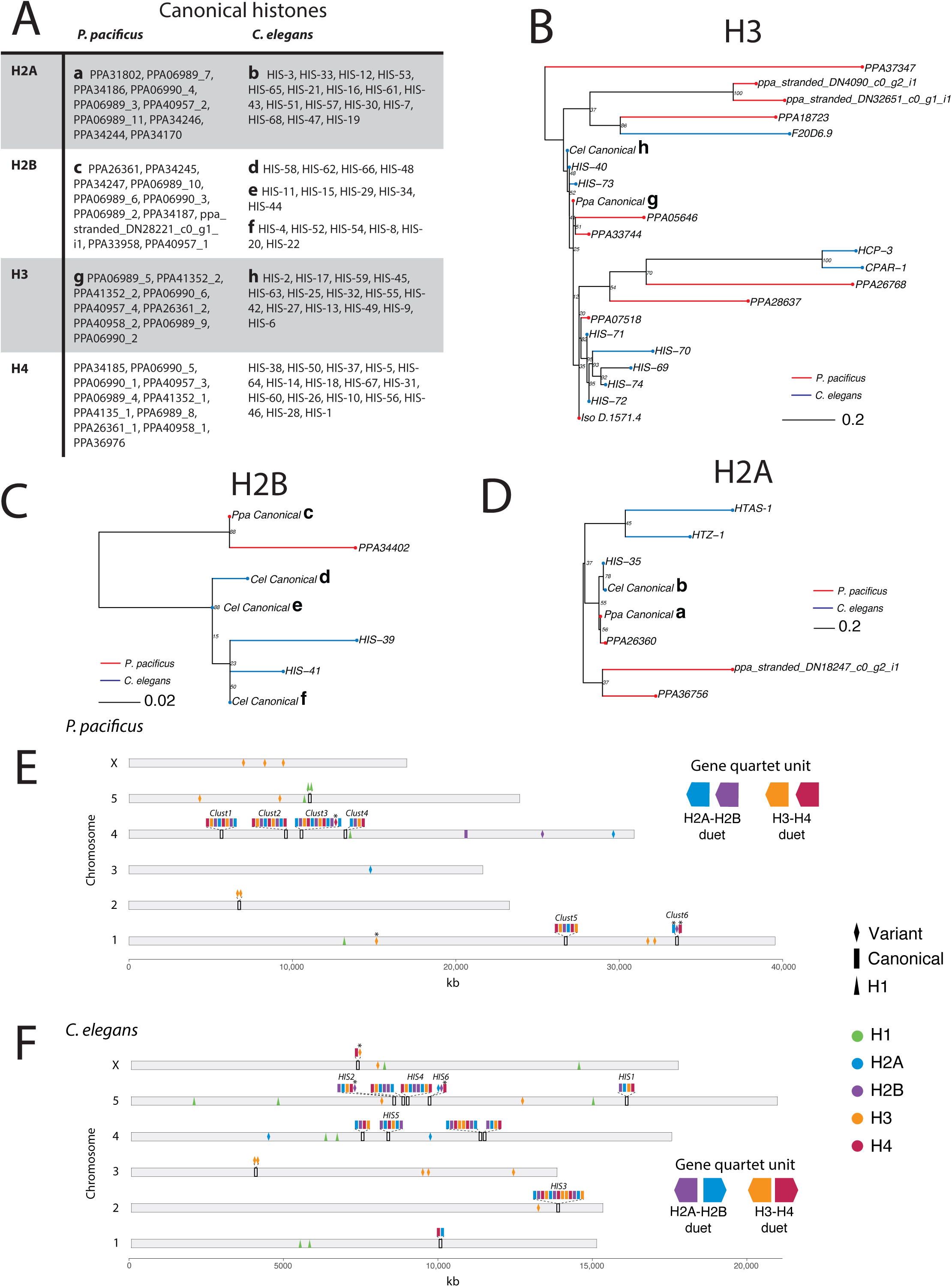
Comparison of *P. pacificus* and *C. elegans* histones. A) Table of *P. pacificus* and *C. elegans* canonical histones. *P. pacificus* gene names ending with an underscore and number were manually reannotated (Additional File 5) B-D) H2A, H2B, and H3 phylogenies generated using maximum likelihood from aligned histone amino acid sequences and midpoint rooted. Branch length reflects the average number of amino acid substitutions per site. Canonical histones indicated with a lowercase letter, corresponding with the table in 2A. Note: *P. pacificus* H4 contains one variant not shown, PPA06989_1 E-F) Histone chromosomal positions. Asterisk (*) indicates cases where a variant is replication-dependent (containing a 3’ UTR hairpin associated with replication dependency) or a canonical gene is not replication-dependent (not containing a 3’ UTR hairpin). *C. elegans* clusters and names as described elsewhere (Pettitt *et al*., 2001; Roberts *et al*., 1987; Roberts *et al*., 1989).

In *C. elegans,* 11 primary histone gene clusters comprised of “quartet” units – tandem repeats of canonical H2B, H2A, H3, and H4 – have been previously characterized (Roberts et al. 1987; Pettitt et al. 2002). Therefore, we hypothesized that histone cluster duplications, deletions, or other genomic rearrangements may explain the differences in the number of canonical histone genes between the two species. Mapping both *P. pacificus* and *C. elegans* histones to their respective chromosomal position revealed that *P. pacificus* contains six canonical histone gene clusters. (Fig. 2E-F). Excluding the replication-independent and largely variant containing *C. elegans* HIS6 cluster, and a homologous *P. pacificus* cluster we called Clust6, each *P. pacificus* cluster is comprised of quartet gene repeats similar to *C. elegans*. However, the two species contain different quartet units: the *P. pacificus* quartet typically contains genes in the order H2A, H2B, H3, and H4 (with an exception in Clust3) all oriented in the same direction (i.e., head-to-tail), whereas in *C. elegans* the histone genes are typically found in the order H2A, H2B, H4, H3 (with an exception in the first chromosome 4 cluster), with H2A and H4 oriented opposite of H2B and H3 (i.e., head-to-head). We found no *C. elegans* clusters containing the *P. pacificus* quartet unit or vice versa. Finally, we asked how conserved this pattern is — do other *Pristionchus* or *Caenorhabditis* species contain similar quartet orders and orientations? We used BLAST to identify histone clusters in *Caenorhabditis bovis, Pristionchus mayeri,* and the non-*Pristionchus* diplogastrid *Allodiplogaster sudhausi* (Supplementary Fig. 3). Within these related species, there is variability in the order of the histone genes within a quartet. However, within each quartet, the H2A-H2B and H3-H4 “duet” orientation is conserved. All *C. bovis* clusters contain H2A-H2B and H3-H4 duets where the genes are in a head-to-head orientation as in *C. elegans*. In contrast, *P. mayeri* and *A. sudhausi* only contain head-to-tail oriented duets as seen in *P. pacificus*. Thus, the order of duets is more evolutionarily labile than the orientation. These data also suggest that there is a deep divergence of the duplication events that formed the *Pristionchus* and *Caenorhabditis* histone gene clusters. We propose that *P. pacificus* and *C. elegans* histone clusters underwent lineage-specific expansions after the duplication of unique duet units. The difference in the number of canonical genes between the two species is likely a result of this independent cluster evolution.

### Characterization of *P. pacificus* histone post-translational modifications, writers, and erasers

We were interested in defining the histone PTMs present in *P. pacificus* to pair with our putative epigenetic gene dataset. Previously, ten histone PTM sites (H3K4me1/3, H3K9me3, H3K27me3, H3K27ac, H3K36me3, H4K5ac, H4K8ac, H4K12ac, and H4K16ac) have been observed using ChIP-seq and/or western blot (Werner et al. 2018; Werner et al. 2023). However, known PTMs are limited to the antibody of choice, and antibody off-target binding can generate misleading interpretations (Grzybowski et al. 2015). To better characterize the suite of histone PTMs in *P. pacificus*, we performed LC-MS/MS on histones extracted from adult worms. Using this more sensitive and comparatively unbiased method, we detected an additional 45 acetylation, methylation, ubiquitination, phosphorylation, propionylation, and hydroxybutyrylation sites on H2B, H3, and H4 (Fig. 3, Supplementary Fig. 4). Histone acetylation, methylation, ubiquitination, and phosphorylation are well characterized across species, however, histone propionylation and hydroxybutyrylation have only recently been identified through targeted mass spectrometry (Kebede et al. 2017; Zhou et al. 2022). We were unable to detect any modifications on H2A in *P. pacificus*. H2A post-translational modifications are less well characterized and thought to be less common than those on their other histone counterparts (Beck et al. 2006). One notable exception is H2A mono-ubiquitination (at lysine K119 in mammals, K120 in *C. elegans*), previously reported in *C. elegans* embryos (Samson et al. 2014). From our data, we cannot conclude whether this mark or any other H2A modifications are necessarily absent in *P. pacificus*: they may be present at levels below our detection threshold or only detectable at different developmental time points. Nevertheless, this data set provides a biochemical inventory of histone PTMs in *P. pacificus* that can be used to complement our bioinformatic inventory of genes.

**Figure 3:**
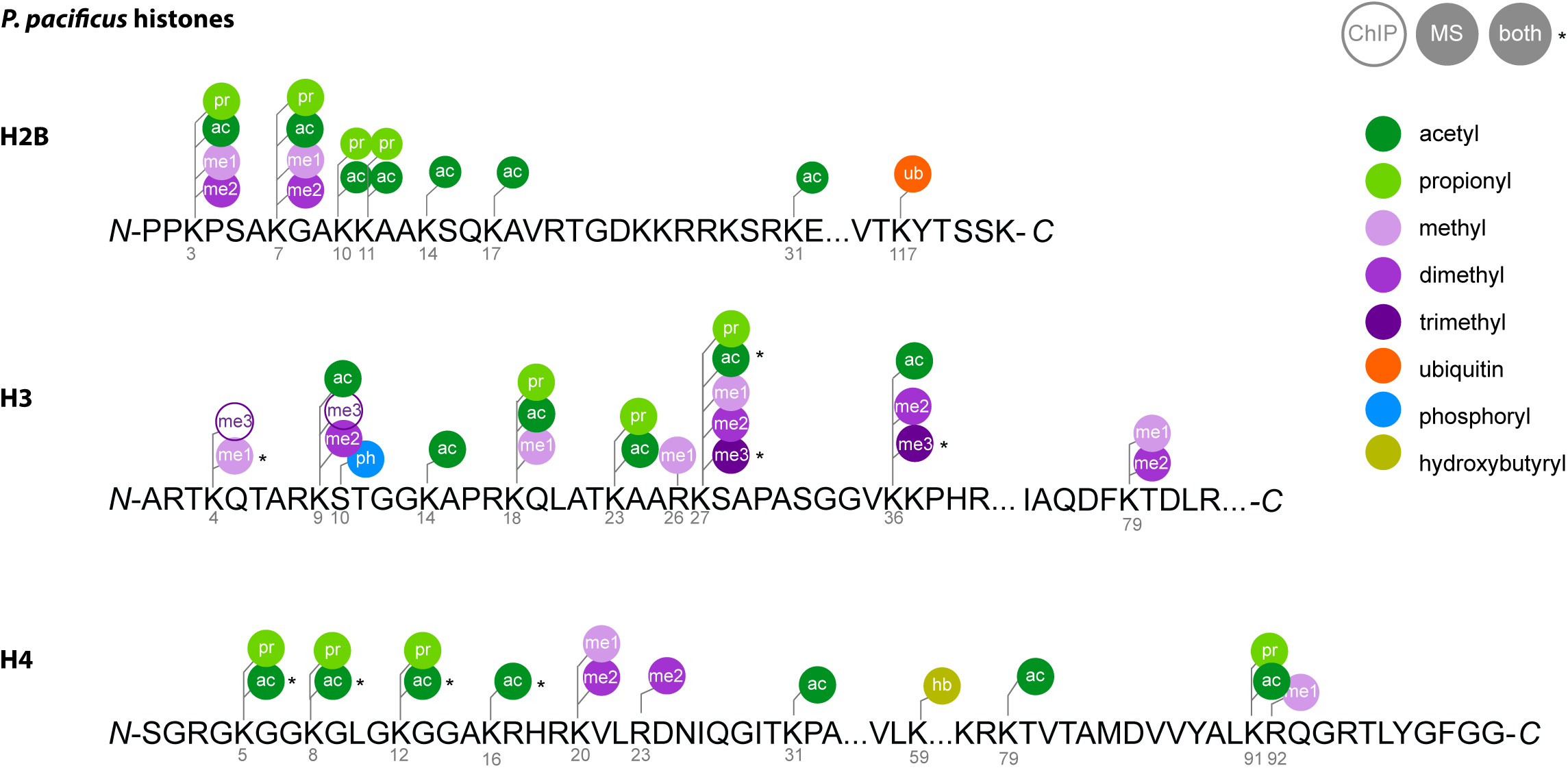
*P. pacificus* histone post-translational modifications. Summary of all PTMs detected in histones extracted from adult *P. pacificus* pellets 72 hours post synchronization. PTMs were detected with LC-MS/MS (PEP score <0.01; this study), ChlP seq (Werner *et al*. 2018; Werner *et al*. 2023), or both.

We then asked whether *P. pacificus* contains orthologs of human and *C. elegans* acetyltransferases, deacetylases, methyltransferases, and demethylases to pair with our histone LC-MS/MS data, as these are the most well-characterized histone-modifying enzymes involved in gene regulation. To this end, we used our orthology dataset and orthogroup phylogenies to predict *P. pacificus* orthologs of human and *C. elegans* enzymes. From this analysis, we could predict 83 *P. pacificus* orthologs of previously characterized human and *C. elegans* histone-modifying enzymes (Table 2-3). Twenty-one of the *C. elegans* or human proteins have known preferred substrates that have been detected in *P. pacificus* either from mass spectrometry (this study) or antibody-based methods (Table 2; Additional File 9; Werner et al. 2018; Werner et al. 2023). In combination, these data establish the repertoire of histone PTMs present in *P. pacificus* and provides candidates for the writers and erasers of specific marks.

**Table 2:**
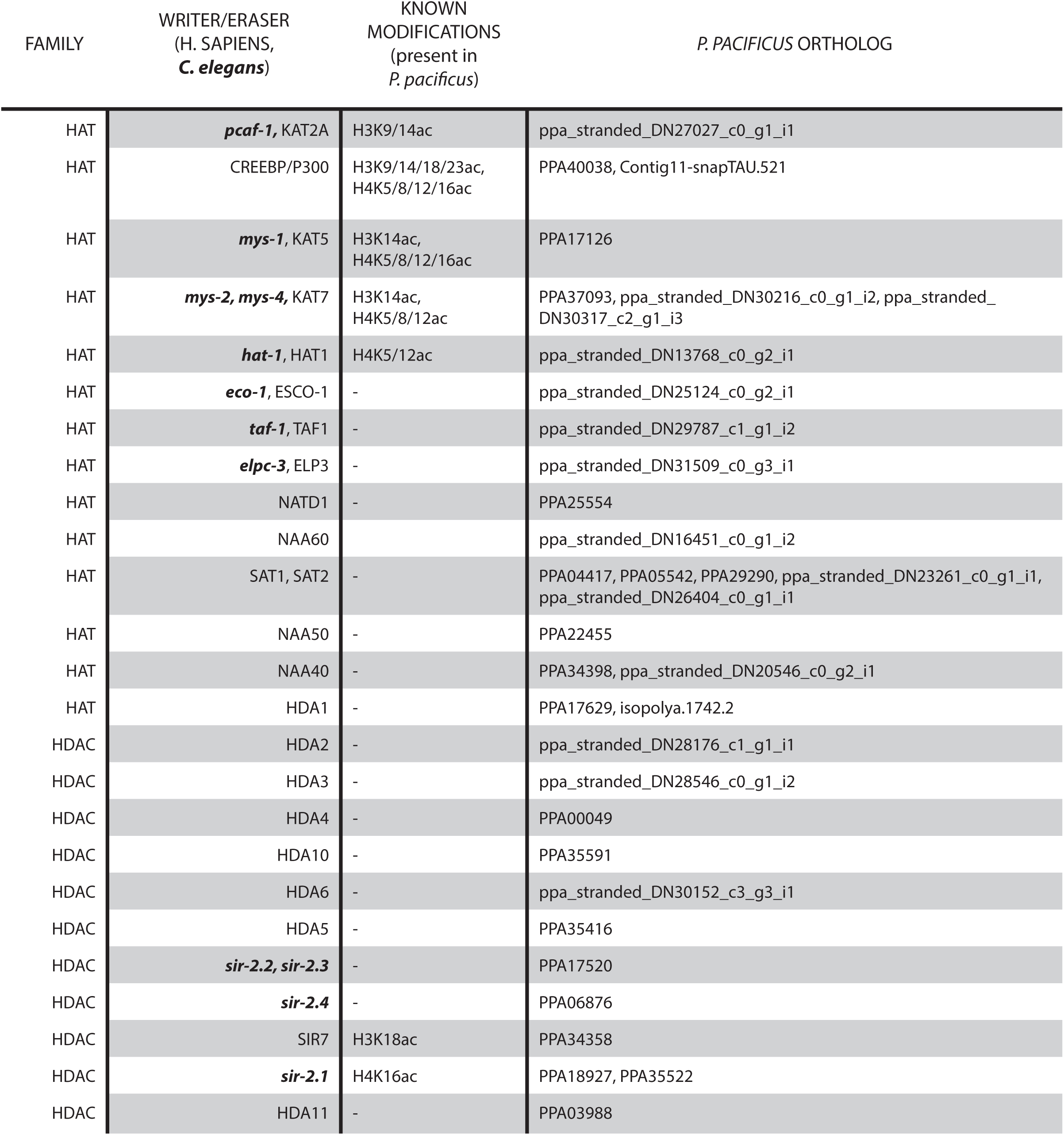
*P. pacificus* histone acetyltransferases and deacetylases. *P. pacificus* orthologs of known human/*C. elegans* HATs and HDACs, identified from orthogroup phylogenetic analysis.

### The methyltransferase complex PRC2 was lost in *P. pacificus* and related *Pristionchus* nematodes

From our “epigenetic toolkits” in *P. pacificus* and *C. elegans,* we noted many instances of species-specific gene gains and losses (Table 1, Supplementary Fig. 1). Of these, we were surprised to find that *P. pacificus* was missing an ortholog of MES-2/EZH2 (*C. elegans* ID/human ID) (Table 3, Fig. 4A). MES-2/EZH2 is the enzymatic component of the highly conserved PRC2 complex, which deposits H3K27me3 to provide maintenance of gene repression (Ahringer and Gasser 2018). The PRC2 complex is the only known eukaryotic H3K27me3 catalyst *in vivo* (A SET protein in the PBCV-1 virus is the only other enzyme currently reported to catalyze this mark; Mujtaba et al. 2008; Wu et al. 2011). Manipulating EZH2 and/or H3K27me3 disrupts cell lineage specification, mammalian X chromosome inactivation and transposon silencing (Yuzyuk et al. 2009; Patel et al. 2012; Walter et al. 2016; Chen and Zhang 2020; Loh and Veenstra 2022). H3K27me3 is also one of the few histone modifications that has been shown to be propagated meiotically and/or mitotically, making it a true epigenetic information carrier (Margueron and Reinberg 2011; Oksuz et al. 2018). In the only eukaryotes for which PRC2 is known to have been lost – the unicellular yeasts *Saccharomyces cerevisiae* and *Schyzosaccharomyces pombe* — the H3K27me3 mark is also absent (Margueron and Reinberg 2011). However, H3K27me3 has been previously detected in *P. pacificus* with ChIP (Werner et al. 2023), and our new analysis confirmed this finding with LC-MS/MS (Fig. 3, Supplementary Fig. 4). The presence of H3K27me3 – but the seeming absence of its writer MES-2 – compelled us to investigate further.

**Figure 4:**
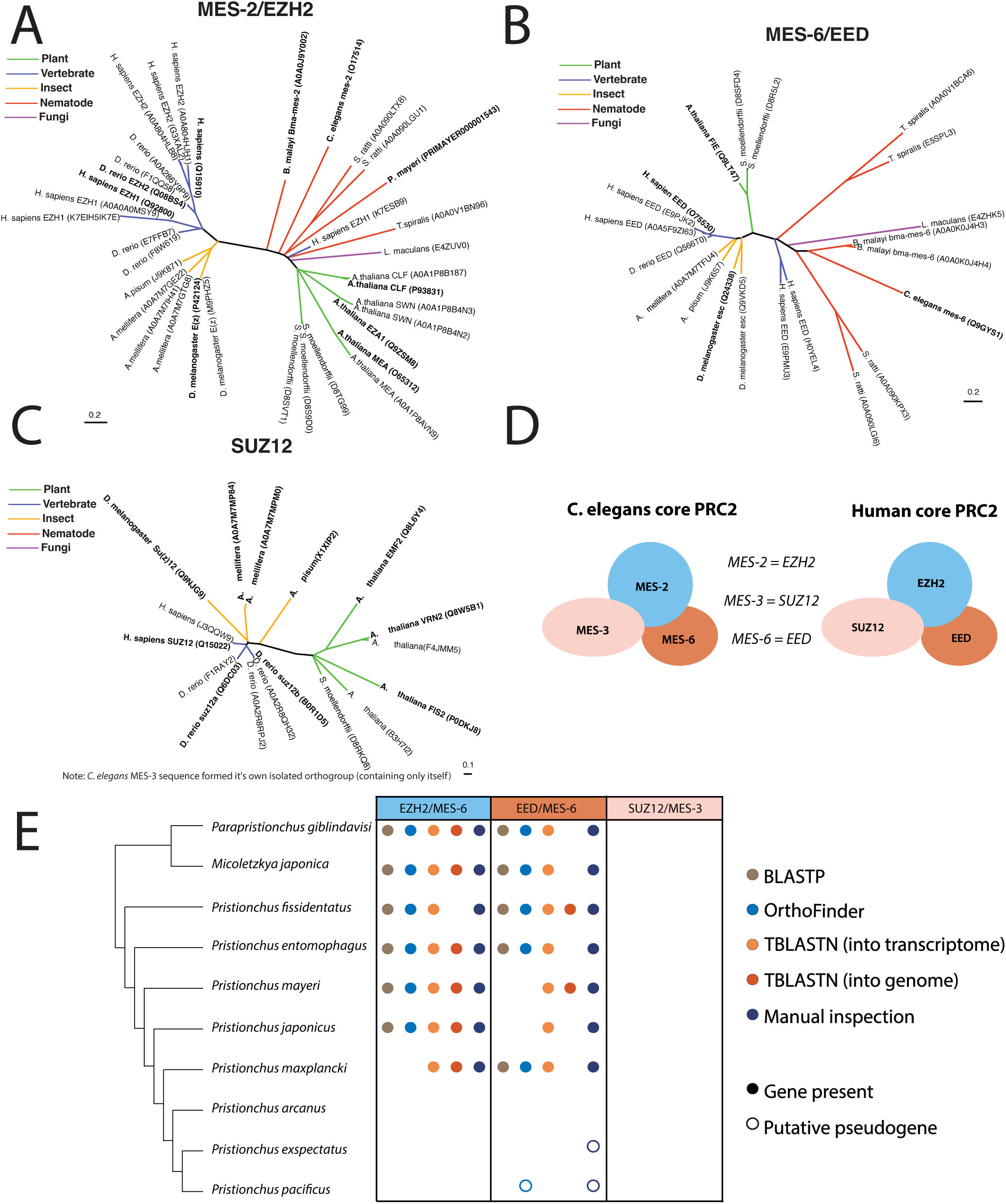
The PRC2 methyltransferase is not conserved in *Pristionchus*. A-C) MES-2/EZH2, MES-6/EED, and MES-3/SUZ12 orthogroups were generated using the pipeline illustrated in Figure 1. UniProt accession numbers are listed in parentheses. For any nearly-identical UniProt entries for the same protein (reflecting either separate protein isoforms or database redundancy), the primary (verified) UniProt entry is in bold. D) Schematic of human and *C. elegans* PRC2 orthologs. E) ldentification of PRC2 component orthologs in the *Pristionchus* phylogeny via BLASTP (e<10E-29); Orthology clustering via OrthoFinder of the10 listed nematodes, *C. elegans*, and humans; TBLASTN into the transcriptome (e<10E-29); TBLASTN into the genome (e<10E-5); and (in cases where the genomic loci of interest could be identified), manual inspection of mapped RNA-seq reads.

**Table 3:** *P. pacificus* histone methyltransferases and demethylases. *P. pacificus* orthologs of known human/*C. elegans* HMTs and HDMTs, identified from orthogroup phylogenetic analysis.

| FAMILY | WRITER/ERASER<br>(H. SAPIENS,<br>*C. elegans*) | KNOWN<br>MODIFICATIONS<br>(present in<br>*P. pacificus*) | *P. PACIFICUS* ORTHOLOG |
| --- | --- | --- | --- |
| HMT | **set-17**, PRDM9 | H4K4me1 | ppa_stranded_DN30240_c0_g14_i5 |
| HMT | **set-30** | H3K4me1 | ppa_stranded_DN28741_c0_g1_i1 |
| HMT | **set-2**, SETD1A/B | H3Kme3, H3K4me1 | PPA00927 |
| HMT | **set-16** | H3K4me3 | ppa_stranded_DN28698_c0_g2_i1 |
| HMT | **met-2**/, SETDB1/2 | H3K9me2, H3K9me3 | ppa_stranded_DN31410_c4_g4_i1, PPA43093 |
| HMT | SUV39H1 | H3K9me3 | PPA41803, ppa_stranded_DN22002_c0_g2_i1 |
| HMT | **set-18** | H3K36me2 | isopolya.8103.2 |
| HMT | **mes-4** | H3K36me2 | PPA11725, PPA27330 |
| HMT | **lin-59**, ASH1L | H3K36me2,<br>H3K36me3 | ppa_stranded_DN31241_c1_g7_i1 |
| HMT | **set-1**, SETD8 | H4K20me1 | PPA02950 |
| HMT | **set-4**, KMT5B | H4K20me2 | PPA04336 |
| HMT | **prmt-5**, PRMT5 | - | ppa_stranded_DN8606_c0_g1_i1 |
| HMT | **prmt-1**, PRMT1/8 | - | Contig232-snapTAU.1, PPA00689, PPA35131, PPA37361, PPA40548,<br>PPA42284, PPA43471, ppa_stranded_DN31419_c0_g5_i2, PPA34076,<br>PPA42322 |
| HMT | **prmt-7**, PRMT7 | - | ppa_stranded_DN25326_c0_g1_i2 |
| HMT | **set-11** | - | PPA23400 |
| HMT | **set-23** | - | ppa_stranded_DN25446_c0_g1_i4 |
| HMT | **met-2**, SETDB1/2 | - | ppa_stranded_DN31410_c4_g4_i1, PPA43093 |
| HMT | **set-29**, SETD4 | - | PPA37266 |
| HMT | **set-14**, SMYD1/2 | - | PPA18466, PPA18468 |
| HMT | **set-27**, SETD3 | - | PPA19833 |
|  | set-3 | - | ppa_stranded_DN27277_c0_g1_i1 |
| HMT | **blmp-1**, PRDM1 | - | PPA04978 |
| HMT | **fib-1** | - | isototal.703.1 |
| HMT | **dot-1.1**, DOT1L | H3K79me1,<br>H3K79me2 | PPA03461, PPA05175, PPA21945, ppa_stranded_DN25525_c0_g1_i2 |
| HDMT | **rbr-2** | H3K4me | ppa_stranded_DN20116_c0_g1_i1 |
| HDMT | **jmjd-2** | H3K9me, H3K36me | PPA01940 |
| HDMT | **utx-1** | - | PPA10459 |
| HDMT | **jmjd-4** | - | ppa_stranded_DN29473_c0_g3_i4 |
| HDMT | **psr-1** | - | PPA42591 |
| HDMT | **jmjc-1** | - | PPA05105 |

Two *P. pacificus* proteins (PPA10329 and PPA06844) cluster with *C. elegans* MES-2 from a phylogeny of SET-domain containing proteins (Supplementary Fig. 1A). However, this branch is supported by a low bootstrap (54), and these proteins are not present in the MES-2 orthogroup phylogeny (Fig. 4A). Moreover, manual inspection revealed that homology to MES-2 was essentially limited to the SET domain. Though the *P. pacificus* genome is considered good quality (BUSCO score of 92.6; Rödelsperger et al. 2017), it is also possible that *mes-2* was integrated into a non-assembled part of the *P. pacificus* genome. To account for this scenario, we also performed tBLASTn of raw PacBio reads for orthologs of MES-2, but could not recover any. Thus, neither our bioinformatic pipeline nor a targeted analysis retrieved the catalytic subunit of the PRC2 complex in *P. pacificus*.

In both humans and *C. elegans,* the PRC2 complex contains three core proteins, each required for catalytic activity: the enzymatic component MES-2/EZH2 and two cofactors MES-3/SUZ12 and MES-6/EED (Fig. 4D) (Jiao and Liu 2015; Ahringer and Gasser 2018; Snel et al. 2022). We first examined our previous orthogroup phylogenies for each of these three proteins to see if we could identify orthologs in *P. pacificus,* as well as in the related *Pristionchus* nematodes *P. exspectatus* and *P. mayeri* (Fig. 4A-C). We only found a *P. pacificus* ortholog of MES-6 that is highly truncated, likely a non-functional pseudogene, and a *P. mayeri* ortholog of MES-2. *C. elegans* MES-3 formed an isolated orthogroup, separate from all other SUZ12 orthologs (consistent with previous reports that MES-3 and SUZ12 are highly diverged orthologs; Snel et al. 2022); neither orthogroup included any *Pristionchus* species. Based on these results, we wondered if PRC2 was lost in multiple *Pristionchus* nematodes.

To further expand our evolutionary analysis of the apareant loss of PRC2 in the *Pristionchus* lineage, we searched for MES-2, MES-6, and MES-3 orthologs in 10 related diplogastrid nematodes which form a ladder-like phylogeny with *P. pacificus* (Prabh et al. 2018), allowing us to connect the presence/absence of PRC2 components to specific branch points. We searched for PRC2 orthologs initially using four different methods: BLASTP, TBLASTN of the genome, TBLASTN of the transcriptome, and OrthoFinder analysis of the 10 diplogastrid nematode proteomes, plus those of *C. elegans,* and humans (Fig. 4E). Both BLASTP and TBLASTN consistently returned slightly more significant results for searches using the human sequences as the search query rather than *C. elegans* (Supplementary Fig. 5A-B). In addition to the previously identified *P. mayeri* MES-2 ortholog, we identified MES-2 and MES-6 orthologs in *P. giblindavisi, M. japonica, P. fissidentatus, P. entomophagus, P. mayeri,* and *P. japonicus.* We also manually curated genome alignments using RNA-seq data (Rödelsperger et al. 2018) to verify gene annotations and search for unannotated orthologs (Fig. 4E). For species with no identifiable *mes-2/mes-6* ortholog, neighboring genes were used to identify the corresponding genomic region. From this analysis, we discovered a putative *mes-6* transcript in *P. exspectatus,* albeit one that contained several stop codons (Fig. 5A). This analysis also revealed several *mes-2* and *mes-6* gene annotation errors in *P. fissidentatus, P. entomophagus,* and *P. japonicus;* re-annotated versions of these genes were used in all further analysis (Supplementary Fig. 6, Additional File 5). No *mes-3* orthologs could be found in any of the diplogastrid examined by any method, indicating that this gene is either absent, or so diverged from the human and *C. elegans* sequences that these methods could not identify it. The combination of all approaches point to *mes-2* and *mes-6* being lost in the last common ancestor of *P. arcanus* and *P. pacificus*.

**Figure 5:**
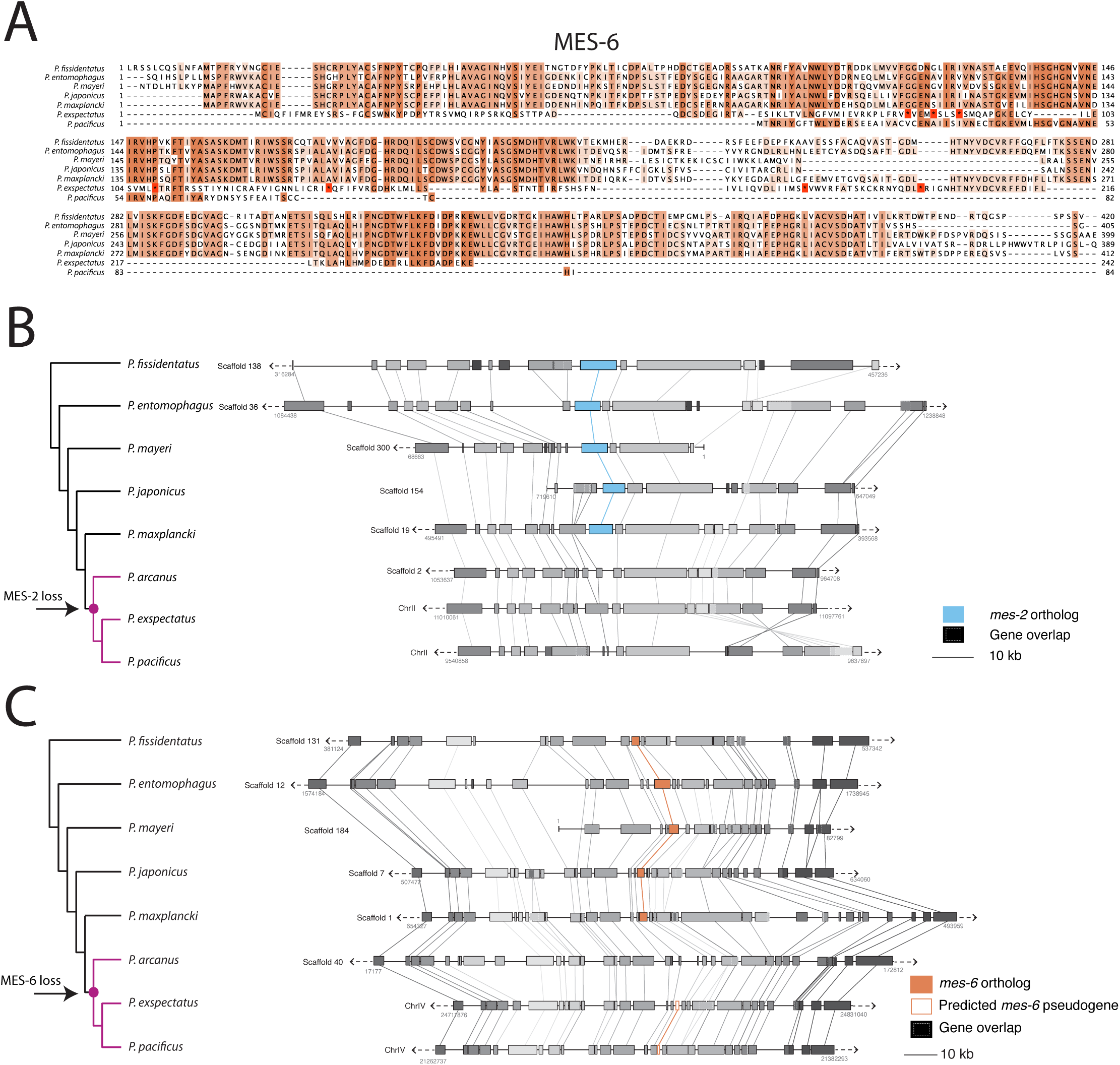
P*r*istionchus *mes-2* and *mes-6* deletion and pseudogenization. A) Protein sequence alignment of MES-6 orthologs in *Pristionchus* nematodes, showing that the *P. exspectatus* ortholog contains many stop codons (*) and additionally departs from the consensus, and the *P. pacificus* ortholog is truncated. B-C) Synteny analysis of the genomic region surrounding *mes-2* and *mes-6* in *Prisitonchus* nematodes. Genes are grouped according to orthogroup membership.

### *Mes-2* and *mes-6* were lost by different evolutionary mechanisms

Pseudogenes often display decreased gene expression and sequence features such as premature stop codons and truncations (Zhang and Gerstein 2004). Truncation of the predicted *mes-6* open-reading frame from *P. pacificus* and multiple stop codons in the predicted *mes-6* transcripts from its sister species *P. exspectatus* indicated pseudogenization (Fig. 5A). However, all *mes-6* genes, including the predicted pseudogenes, displayed expression by RNA-seq (Supplementary Fig. 7B). Premature stop codons or gene truncations were not observed in orthologs of *mes-2*: we either found a presumably functional full-length coding sequence or complete absence. Additionaly, all existing *mes-2* orthologs exhibited expression by RNA-seq (Supplementary Fig. 7A). Thus, at least at this time, there is no apparent signature of decreased expression of *mes-2* along the *Pristionchus* phylogeny that predated its loss in some lineages.

The absence of *mes-2* orthologs (including pseudogenes) could be explained by rapid sequence evolution rendering pseudogenized *mes-2* orthologs unrecognizable, or it could be that the locus has been lost from the genome entirely. We asked whether we could find any indication of large genomic rearrangements that could be responsible for the sudden absence of *mes-2* orthologs. To this end, we analyzed the local synteny of the genomic region surrounding *mes-2* across the *Pristionchus* phylogeny (Fig. 5B). From this analysis, we determined that the local synteny of the genomic region surrounding *mes-2* is conserved across the phylogeny: even though *mes-2* is absent in *P. pacificus, P. exspectatus* and *P. arcanus,* the positions of neighboring genes are relatively fixed. Additionally, the close proximity of the neighboring genes suggests that the *mes-2*-containing region may have been lost entirely in the common ancestor of *P. pacificus* and *P. arcanus*. We also repeated this analysis for *mes-6* and similarly found that the structure of this local region is also well conserved across these nematodes, with pseudogenes replacing functional *mes-6* in *P. exspectatus* and *P. pacificus* (Fig. 5C). Finally, we asked whether there were any repetitive sequences indicative of transposon activity (which have been previously annotated in *P. pacificus* (Athanasouli and Rödelsperger 2022)) that could explain *mes-2’s* loss. However, we did not detect any of these sequences at the *mes-2* locus in *P. pacificus.* Taken together, functional *mes-2* and *mes-6* genes appear to have been lost in the common ancestor of *P. arcanus* and *P. pacificus* but by different evolutionary mechanisms*. Mes-2* appears to have been lost via a specific deletion event, while *mes-6* has undergone a pseudogenization process, leaving behind gene remnants in *P. pacificus* and *P. exspectatus*.

### *P. pacificus* retains H3K27me3 in both its germline and somatic tissue

In *C. elegans,* H3K27me3 is present in essentially all body tissues, yet PRC2 is only essential for maintaining germline H3K27me3 (Bender et al. 2004). H3K27me3 in the *C. elegans* germline is enriched on the X chromosome and is responsible for maintaining X chromosome repression during germline development; loss of any PRC2 component causes loss of all germline H3K27me3 and a maternal effect sterile phenotype (Capowski et al. 1991; Bender et al. 2004; Strome et al. 2014). Because *P. pacificus* appears to have lost PRC2, we asked whether it has correspondingly lost H3K27me3 in its germline but not somatic tissue. In this scenario, the lack of PRC2 in *P. pacificus* would indicate species-specific differences in germline maintenance and explain the detection of H3K27me3 as present excusively in the soma.

We dissected the gonad and intestine (somatic tissue) from wild type adult *P. pacificus* and *C. elegans* worms and stained them with antibodies specific for H3K27me3, H3, and with DAPI to stain nuclei (Fig. 6, Supplementary Fig. 8). As expected, *C. elegans* shows robust H3K27me3 staining in both the gonad and intestine. Interestingly, *P. pacificus* also shows H3K27me3 staining in both tissues. Furthermore, H3K27me3 staining in the *P. pacificus* gonad appears more punctate than the H3 control (Supplementary Fig. 8F). Therefore, H3K27me3 is retained in the *P. pacificus* gonad, potentially for X chromosome silencing (as in *C. elegans*), despite the loss of PRC2.

**Figure 6:**
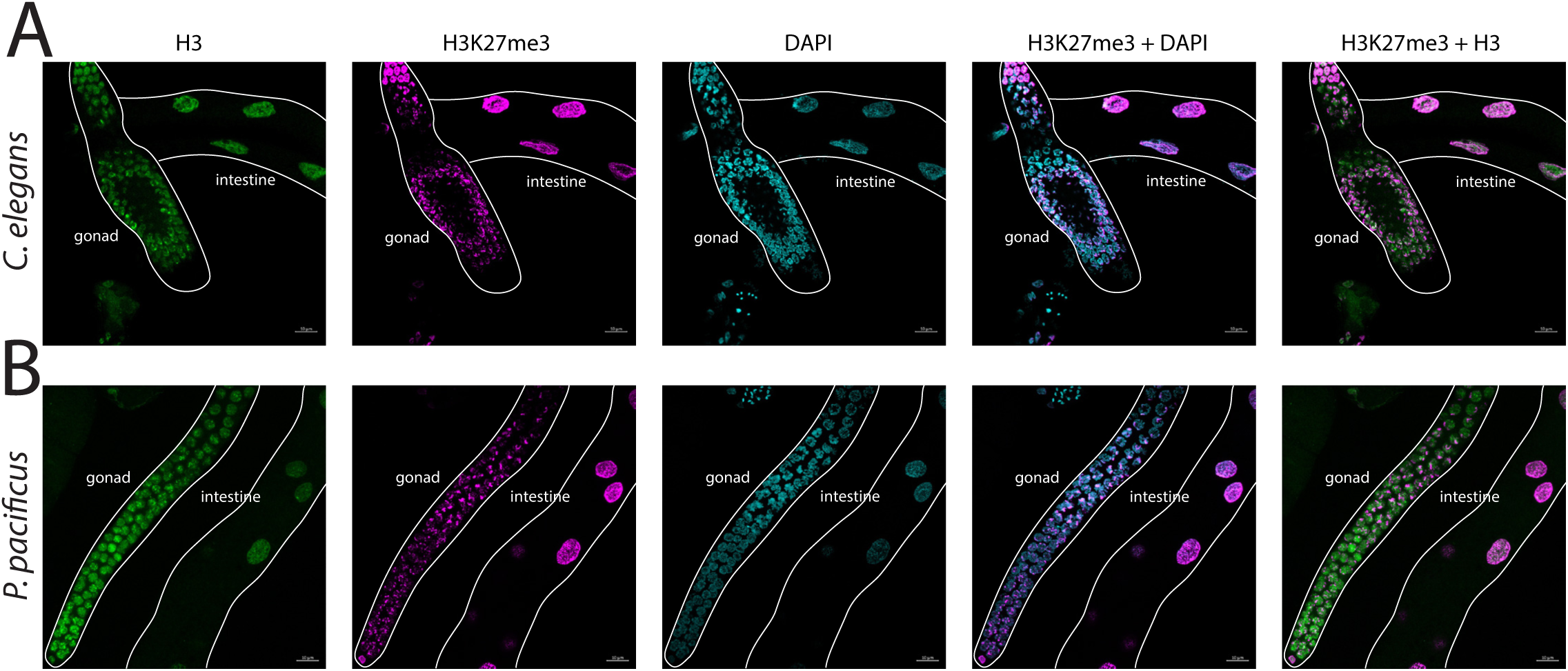
*P. pacificus* contains H3K27me3 in both the soma and the gonad. lmmunofluorescence staining of dissected (A) *C. elegans* and (B) *P. pacificus* gonad and intestine.

## Discussion

Recent pharmacological experiments indicate that epigenetic information carriers, particularly histone PTMs, regulate mouth-form plasticity in the evo-devo model system *P. pacificus* (Werner et al. 2023). However, validating this result and connecting it to the action of individual genes requires functional information on the repertoire of epigenetic writers and erasers encoded within the *P. pacificus* genome. To address this shortcoming, we have created an inventory of putative epigenetic genes in *P. pacificus,* as well as in *C. elegans,* using a domain and orthology-informed pipeline. These datasets provide a foundation for experimental analysis of plasticity and gene regulation in both *P. pacificus* and *C. elegans*. Additionally, a comparison of the epigenetic toolkits between both model nematodes highlights unexpected facets of evolutionary divergence.

First, there was a significant difference in histone gene composition. In both species, core “quartet units” of canonical histone genes have multiplied throughout the genome to form clusters (Roberts et al. 1987; Pettitt et al. 2002). Despite the high conservation of histone genes themselves, the *P. pacificus* and *C. elegans* histone clusters contain quartets comprised of distinct duets (in terms of gene order and orientations), with no instances of the *P. pacificus* duets being found in *C. elegans* or vice versa. We also found that the respective *C. elegans* and *P. pacificus* histone gene duets are conserved among each’s close relatives (in *C. bovis,* for *C. elegans;* and in *P. mayeri* and *A. sudhausi,* for *P. pacificus*). These findings are consistent with a deep divergence of *P. pacificus* and *C. elegans* (estimated at 80-200 million years; Howard et al. 2022). We propose that the observed differences in the total canonical histone genes between *P. pacificus* and *C. elegans* are due to independent histone gene cluster formation — indicated by the duplication of distinct gene duets in each species. Going forward, it will be interesting to see when each histone gene cluster arose, which may help to resolve relative nemstode phylogenetic positions. Recent progress in nematode genomics is providing the raw data needed for such analyses (Prabh et al. 2018; Rödelsperger 2021; Wighard et al. 2022).

While we saw the greatest variation between *P. pacificus* and *C. elegans* in terms of histone gene count, our results also reveal differences in the types of histone-modifying enzymes present. In particular, we were unable to identify *P. pacificus* orthologs of the PRC2 components MES-2 and MES-3 despite searching with several distinct methods: orthology clustering (OrthoFinder), BLASTP, TBLASTN into the genome, TBLASTN into the transcriptome, and manual inspection of RNA-seq reads mapped to the genomic locus. We did identify a *mes-6* pseudogene in *P. pacificus*, providing further evidence of a non-functional *P. pacificus* PRC2 since MES-6, MES-2, and MES-3 are each required for PRC2 catalytic function (Jiao and Liu 2015; Ahringer and Gasser 2018; Snel et al. 2022). It is formally possible that *mes-2* and *mes-6* genes duplicated, were integrated into a non-assembled part of the genome, and their respective parental genes pseudogenized. However, we should still be able to recover tBLASTn hits from the raw sequencing reads, and could not. Moreover, missing genes should appear randomly across the phylogeny and should also be random in regard to the presence/absence of constituent members of complexes. The phylogenetic signature of a single loss, and the fact that no intact members of the complex are recoverable within a phylogenetically well-defined cluster of closely-related species argues against this possibility. Thus, although we were intitially skeptical that the PRC2 complex was absent in *P. pacificus*, we ultimately could not refute this interpretation with multiple independent methods. We conclude that the PRC2 complex was lost in the last common ancestor of *P. pacificus* and *P. arcanus*.

Evolutionary changes in the composition and functionality of PRC2 complex proteins have been documented between invertebrates and humans. For example, the main SET methyltransferase protein was duplicated in vertebrate evolution, leading to two genes: EZH1 and EZH2. EZH2 is most similar to invertebrate orthologs and is responsible for the majority of methyltransferase activity in mammels, therefore we focused our analyses on this gene (Fischer et al. 2022). However, to our knowledge, the complete loss of PRC2 has yet to be documented for any multicellular organism. This raises the question of how the loss of PRC2 is tolerated in *P. pacificus*. The few eukaryotes without PRC2 (the single-celled yeasts *S. pombe* and *S. cerevisiae*) have simultaneously lost H3K27me3 (Shaver et al. 2010). However, we confirmed using LC-MS/MS that H3K27me3 is present in *P. pacificus*. This finding suggests that an enzyme other than PRC2 must catalyze H3K27me3 in *P. pacificus*.

In *C. elegans*, PRC2 is primarily expressed in the germline, though with some expression recently reported in the endoderm and mesoderm (Bender et al. 2004; Engert et al. 2018; Vaart et al. 2020). Knocking out *mes-2* leads to a maternal effect sterile phenotype and reduces H3K27me3 in the germline. However, in these experiments, H3K27me3 is not lost in the soma, indicating that in *C. elegans,* MES-2 is responsible for maintaining germline but not somatic H3K27me3 (Bender et al. 2004). Presumably, another *C. elegans* H3K27me3 methyltransferase is responsible for maintaining somatic H3K27me3, though to date, the identity of this somatic writer is unknown. Our immunostaining experiments in *P. pacificus* demonstrate that, despite the absence of PRC2, H3K27me3 is still maintained in both the gonad and somatic tissue. One possibility is that both *P. pacificus* and *C. elegans* share an uncharacterized histone methyltransferase, responsible for somatic H3K27me3 in *C. elegans* and global H3K27me3 in *P. pacificus.* Of course, these functions and their enzymatic writers may be unrelated. Lastly, H3K27me3 is highly enriched over the super-gene locus controlling mouth-form in *P. pacificus* (Werner et al. 2023). Thus, identifying the enzyme responsible for H3K27me3 in *P. pacificus* may expand our understanding of plasticity and the evolution of epigenetic mechanisms.

## Acknowledgements

We thank past and present members of the Werner lab for all input and discussions about this project, especially Madelyn Purnell for the many discussions about gene evolution. We also thank Ralph Sommer for all guidance and advice. This work was supported by the National Institute of General Medical Sciences (R35GM150720) and the School of Biological Sciences at the University of Utah. A.B. was also supported by an NIH funded training grant (T32-GM122740) and the National Science Foundation Graduate Research Fellowship (DGE-2039655)

## Data availability

The LC/MS-MS has been deposited to the ProteomeXchange Consortium via the PRIDE partner repository with the dataset identifier PXD046748. The code and data used to create the histone position and synteny plots can be found at https://github.com/audreybrown1/Brown-et-al.-2023-library. All other data needed to evaluate the conclusions in this paper is included as supplementary files, which are referenced throughout the text. Additional File 1 lists the reference Pfam domains and Additional File 2 contains the reference model organism epigenetic genes used in the domain and orthology informed pipeline. Additional File 2 contains the InterProScan results for *P. pacificus.* Additional File 4 contains the initial OrthoFinder Orthogroup results. Additional File 5 contains gene position information for any manually reannotated genes. Additional File 6 contains our predited putative epigenetic genes for *P. pacificus.* Additional File 7 contains the InterProScan results for *C. elegans* and Additonal File 8 contains our predicted putative epigenetic genes for *C. elegans.* Additional File 9 contains all references on histone modifying enzymes used to generate Tables 2-3. Additional File 10 contains the OrthoFinder Orthogroup results using *Pristionchus* species. Additional Files 11-12 contain gene position data used to generate the *mes-2* and *mes-6* synteny plots (Fig. 5B-C). Additional Files 13-14 contain the histone gene position information used to generate the histone position plots (Fig. 2E-F).

## Author contributions

A.B., A.B.M., C.J.W., and M.S.W. designed the bioinformatic analysis pipeline; A.B. and A.B.M. performed bioinformatic identification of epigenetic genes; A.B. and M.W. analyzed histone gene cluster conservation; A.B. performed all analyses on PRC2 conservation; M.F.W. performed LC-MS/MS with guidance from B.M. A.B. created all phylogenies with guidance from C.J.W. A.B. and S.G. performed immunofluorescence experiments with guidance from O.R. Writing was by A.B. and M.S.W. with input from all authors.

